# Real-world results from a Machine Learning-guided, phenotypic High-Throughput Screen for novel antibiotics

**DOI:** 10.64898/2026.06.22.733866

**Authors:** Paul Lukacs, Kevin C. Hare, Shilpa George, Graham Hone, Gayatri Gollapudi, Lisa Wang Jarantow, Jenna Pellegrino, Alita Miller, Kurt Thorn

## Abstract

Antimicrobial resistance is an urgent global health threat, with over 2.8 million multidrug-resistant infections killing over 35,000 annually in the US. Machine Learning (ML) has emerged as a potential solution to improve efficiency of antibiotic high-throughput screens (HTS). We report ML-guided high-throughput screening against *E. coli*. Large-scale Learning-to-Rank models were trained on public and proprietary datasets to maximize phenotypic inhibition and minimize human cell cytotoxicity. We evaluated several pre-plated compound libraries and a set of “cherry-picked”, structurally novel compounds. We screened against a hyperpermeable lptD^-^ mutant, followed by hit confirmation, profiling, cytotoxicity counter-screening, and MOA determination. Results demonstrated a doubled hit rate and 3X fewer toxic hits. Additionally, activity improved against both Wild Type *E. coli* and the lptD^-^ mutant. ML models showed robust predictive power on structurally dissimilar compounds. The combination of large-scale HTS, ML innovation, and both library-wise selection and cherry-picking strategies distinguishes this study in the antibiotic discovery field.

## INTRODUCTION

Antimicrobial resistance represents one of the most urgent health threats of the 21^st^ century. Multidrug-resistant bacterial infections result in millions of hospital days and billions in healthcare costs annually (Diéguez-Santana & González-Díaz, 2023). The situation has been exacerbated by the pharmaceutical industry’s retreat from antibiotic development towards more lucrative chronic treatments (Durrant & Amaro, 2015). This has created a significant innovation gap, with few novel antibiotics entering the market despite the escalating crisis (Balakrishnan, 2022; Brown & Wright, 2016).

Small molecules have dominated approved and pending antibiotics (Silver, 2011, Butler et al., 2017). Historically, pharmaceutical companies have relied on HTS to identify potential antibiotic leads with new MOAs. Unfortunately, these efforts have proven inefficient. For example, between 1995 and 2001, GlaxoSmithKline conducted 70 HTS campaigns yielding only five leads — a success rate four to five times lower than for other drug targets (Durrant & Amaro, 2015; Payne et al., 2007). The challenges are multifaceted: for example, small molecules must not only demonstrate activity against bacterial targets but also overcome barriers like bacterial membranes (particularly in Gram-negative species) and efflux pumps (Silver, 2011); additionally, traditional HTS is prohibitively expensive for academic research groups, who are increasingly important players as pharmaceutical companies have shifted focus (Tommasi et al., 2015).

Machine Learning (ML) has emerged as a potential solution to alleviate these problems by:

1. Improving hit rates: ML can prioritize compounds likely to demonstrate favorable properties (Jukič & Bren, 2022).
2. Efficiently exploring chemical space: computational methods can generate predictions across chemical spaces orders of magnitude more rapidly than compounds can be screened in the laboratory or assessed by a medicinal chemist (G. Liu & Stokes, 2022).

In this study, we focus on so-called enumerative ML workflows as opposed to generating virtual compounds *de novo*. The enumerated compounds can come from commercial or virtual libraries or can be combinatorially created based on synthesis schemes. We focus on enumerative methods for two reasons: first, they resemble conventional HTS and virtual screening more closely and are more compatible with real-world hit finding pipelines (Vamathevan et al., 2019); second, generative models are known to propose unrealistic or difficult-to-synthesize compounds (Gao & Coley, 2020; Stanley & Segler, 2023). ML-guided HTS is analogous to virtual screening with two main differences: first, the computationally expensive predictor such as molecular docking is replaced by a more scalable, data-driven ML model (E. Graff et al., 2021); second, ML models can predict phenotypic activity that is beneficial in antibacterial drug discovery when Gram-negative bacterial efflux and membrane permeation confound protein binding (Nikaido, 1994), or when the target protein is unknown (Childers et al., 2020; Croston, 2017).

ML-guided HTS studies fall into two categories: retrospective and prospective. Retrospective studies benchmark ML guidance through cross validation using literature data. Conversely, prospective studies acquire previously unseen, ML-prioritized compounds, test them experimentally, then evaluate the improvement due to ML guidance (the “ML lift”). Retrospective studies are useful as a low-cost, first-line approach to estimate the benefits of ML guidance – many studies report promising performance in a variety of use cases (Castillo-Garit et al., 2015; Cherkasov, 2005; Dalecki et al., 2019; Karakoc et al., 2006; Marrero-Ponce et al., 2005; Masalha et al., 2018; Yang et al., 2009). Importantly, retrospective studies can overestimate performance due to well-known biases and do not yield insight into operational aspects of ML-guided drug discovery (van Giffen et al., 2022).

Prospective studies better emulate real-world hit finding. Several studies explored ML-guided screening in academic settings and reported impressive ML lift (Cesaro et al., 2023; Ivanenkov et al., 2019; Murcia-Soler et al., 2004; Rahman et al., 2022; Stokes et al., 2020; Wang et al., 2014; Wong et al., 2024). The details of these studies are summarized in Table 1. To our knowledge, Scalia et al. reported the only study to date that evaluated ML guidance for antibacterial discovery in a non-academic setting. They used GNEprop, a Deep Learning model trained on an extensive in-house dataset of 1.2 million compounds with measured activity against two *E. coli* strains. The top-scoring 345 of over 1.7 billion compounds were selected and a hit rate of 24% was observed, a 90-fold improvement over the fraction of actives in their training set. They noted that several of the found actives were dissimilar to training compounds (Scalia et al., 2024).

**Table 1.** Summary of prospective ML-guided antibiotics screening studies.

| Reference | Application | ML technique | Training data size | Compound selection | Result |
| --- | --- | --- | --- | --- | --- |
| Murcia-Soler et al., 2004 | *S. aureus*<br>*E. faecalis* | Multilayer Perceptron | 433 |  | 4 experimentally validated hits |
| Wang et al., 2014 | *S. aureus* | Various | 4,541 | Top 56 of 7,500 | 12 were active; Naïve Bayes classifier best model |
| Ivanenkov et al., 2019 | *E. coli* | Various | 140,000 | 5,000 | Identified several leads; Kohonen Map best model |
| Stokes et al., 2020 | *E. coli* | Directed Message Passing Neural Network (DMPNN) | 2,315 | ~5,000* | Found Halicin, a promising broad-spectrum antibiotic |
| Rahman et al., 2022 | *B. cenocepacia* | DMPNN | 29,537 | Top ~120 of ~225,000** | At least 14-fold lift; broad-spectrum hits |
| Cesaro and Fuente-Nunez, 2023 | *A. baumannii* | DMPNN | 7,688 | Top 240 of ~5,000 | 17% hit rate; Abaucin, a hit active against multiple strains |
| Wong et al., 2024 | *S. aureus* | DMPNN | 39,312 | 283 tested | Best compounds associated with growth inhibition and low human toxicity |

**Table 2.**
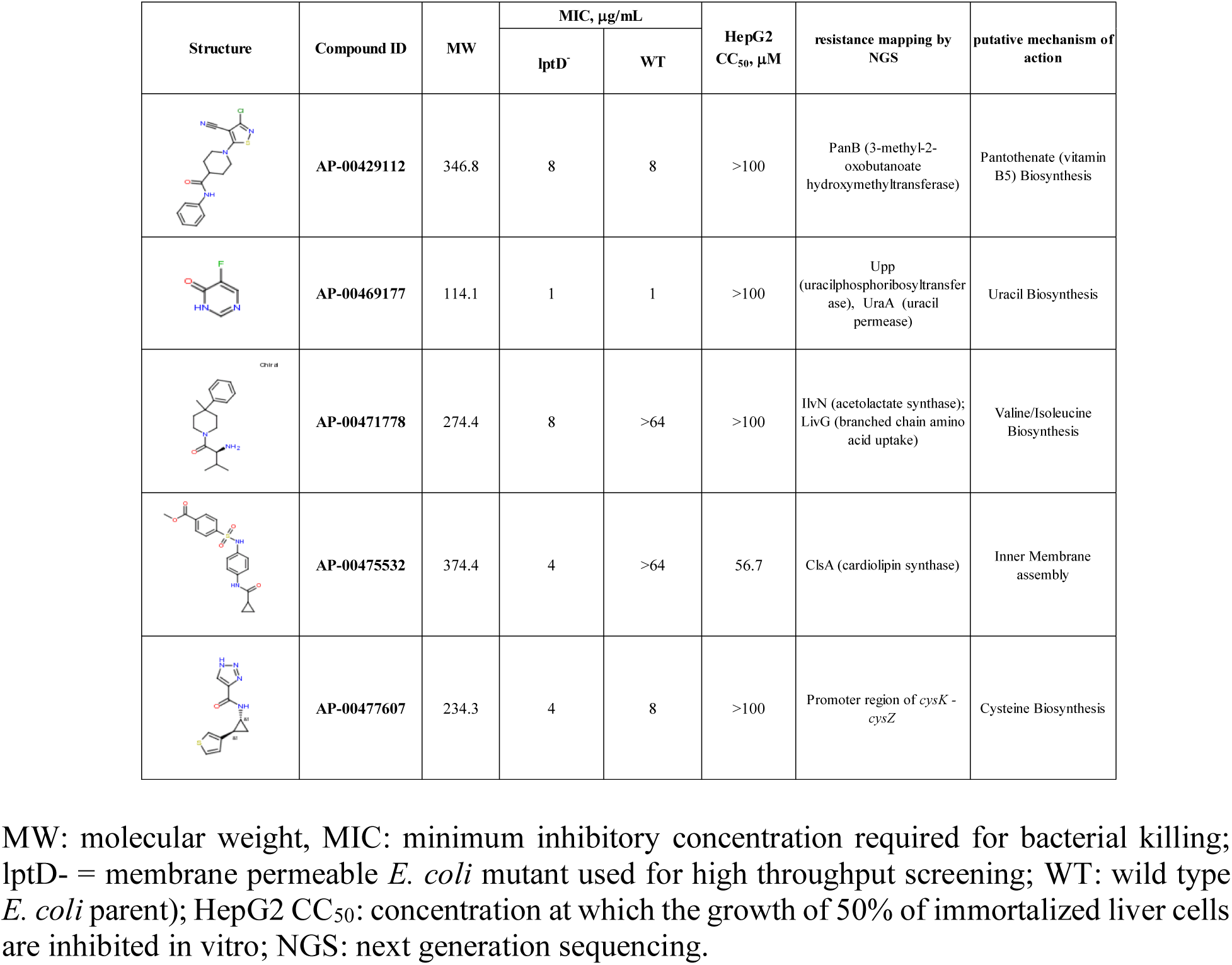
Biological properties of representative hits.

| Structure | Compound ID | MW | MIC, µg/mL |  | HepG2<br>CC<sub>50</sub>, µM | resistance mapping by<br>NGS | putative mechanism of<br>action |
| --- | --- | --- | --- | --- | --- | --- | --- |
|  |  |  | lptD<sup>-</sup> | WT |  |  |  |
|  | AP-00429112 | 346.8 | 8 | 8 | >100 | PanB (3-methyl-2-oxobutanoate hydroxymethyltransferase) | Pantothenate (vitamin B5) Biosynthesis |
|  | AP-00469177 | 114.1 | 1 | 1 | >100 | Upp (uracilphosphoribosyltransferase), UraA (uracil permease) | Uracil Biosynthesis |
|  | AP-00471778 | 274.4 | 8 | >64 | >100 | IlvN (acetolactate synthase); LivG (branched chain amino acid uptake) | Valine/Isoleucine Biosynthesis |
|  | AP-00475532 | 374.4 | 4 | >64 | 56.7 | ClsA (cardiolipin synthase) | Inner Membrane assembly |
|  | AP-00477607 | 234.3 | 4 | 8 | >100 | Promoter region of *cysK* - *cysZ* | Cysteine Biosynthesis |

MW: molecular weight, MIC: minimum inhibitory concentration required for bacterial killing; lptD<sup>-</sup> = membrane permeable *E. coli* mutant used for high throughput screening; WT: wild type *E. coli* parent); HepG2 CC<sub>50</sub>: concentration at which the growth of 50% of immortalized liver cells are inhibited in vitro; NGS: next generation sequencing.

Our study reports real-world results from an ML-guided HTS against a membrane-compromised strain of *E. coli*. We trained Learning-to-Rank (L2R) ML models that integrated public and proprietary datasets to prioritize unseen compounds in terms of their phenotypic inhibition against *E. coli* and human cell cytotoxicity. To evaluate the ability of ML guidance to select compound collections, several pre-plated compound libraries were acquired and screened. In addition, a “cherry-picked” (CP) set of compounds was selected that maximized predicted *E. coli* activity, minimized predicted cytotoxicity and excluded compounds structurally similar to known antibiotics. The acquired compounds were screened against a hyperpermeable lptD^-^ mutant of *E. coli*. Hits underwent hit confirmation, hit profiling, cytotoxicity counter-screening and MOA determination. The HTS results clearly demonstrated hit enrichment and reduction of cytotoxicity; furthermore, hit quality and activity against Wild Type (WT) *E. coli* improved. Notably, the L2R models showed robust predictive power on the unseen, structurally dissimilar screening compounds, both in terms of CP performance and evaluating compound libraries. In our view, this study is unique for the following reasons:

1. the scale and commercial environment of the HTS,
2. the size of the training datasets,
3. the ML formulation novel to antibiotics discovery,
4. compound selection for competing objectives,
5. both library-wise selection and “cherry-picking” individual compounds.

## RESULTS AND DISCUSSION

Approximately 400,000 compounds from Evotec’s screening library were screened for growth inhibition of lptD^-^ *E. coli* in minimal media. The hits were confirmed by retesting in triplicate; confirmed hits were profiled and counter-screened for human cell cytotoxicity. These HTS data, along with public datasets, formed the training set of a set of ML models (predicting antibiotic activity and human cell cytotoxicity) that were used to select compounds in consequent stages of the HTS. The 400,000 compounds were selected without ML guidance, based on human-assessed medicinal chemistry properties. For the balance of this paper, we refer to this screen as the “unguided” baseline. The experimental details of the HTS and technical aspects of the ML models and training data can be found in the Experimental Section.

To estimate expected ranking performance, the trained models were evaluated using cross validation. We held out test sets of 10,000 compounds from relevant subsets of the training sets; subsets were defined based on the data source, i.e., the originating database and assay. In addition, to better evaluate predictive performance on unseen compounds, observations that shared Murcko-Bemis scaffolds with the test observations were removed from the remaining training data. Figure 1 shows the results. Estimated performance is reported in terms of the Receiver Operating Characteristic (ROC) curves and estimated lifts. When the predicted properties are binary classes (active vs. inactive, cytotoxic vs. non-cytotoxic, as in this case), the ROC curves are equivalent to discovery curves, i.e., the horizontal axes express the fraction of library screened in the order of decreasing predicted compound utility, and the vertical axes indicate the fraction of positives (active or non-cytotoxic) discovered at that level of completion. For reference, a diagonal line that represents no enrichment (screening in a random order) is drawn. At any % library screened, dividing the fraction of positives found by the baseline (here, the random, unguided performance) yields the lift at that % library screened: the lifts quantify enrichment in positives due to screening in preferential order based on ML guidance; e.g., a lift of 2 at 1% library screened means that twice as many positives are expected to be found compared to the random baseline. As seen, significant enrichment was estimated in every test set and at every % library screened. The lifts were estimated between 2 and 20, indicating significant benefits from ML guidance. When interpreting the results of this analysis note that 1, the random baseline is uncertain and the confidence bounds are a function of the library size and the number of positives; 2, lift typically decreases with % library screened, indicating diminishing returns as the screening regime leaves the regime of “elite selection”; 3, this analysis assumes single-objective optimization, i.e., selecting for activity or non-cytotoxicity but not for both. Furthermore, we note that cross validation is a backward-looking performance estimation method that can be biased by the train-test splitting scheme.

**Figure 1.**
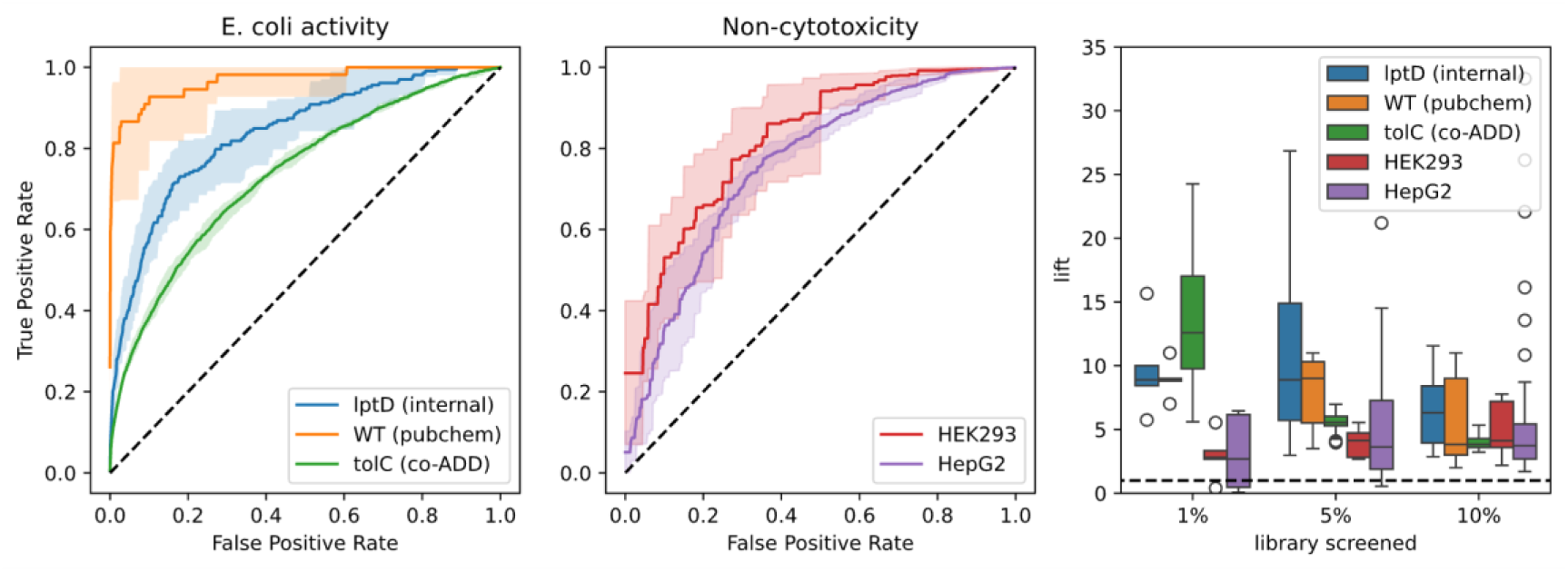
Expected performance of the activity and cytotoxicity rankers estimated in back-testing. Left: *E. coli* activity ROCs when ranking three scaffold-split test sets. Middle: Non-cytotoxicity ROCs when ranking two scaffold-split test sets. The dashed black lines indicate the expectation of a random baseline. Right: estimated lifts as a function of % of library screened for the five test sets. A lift of 2 at 1% screened means that when screening the top-ranked 1% of a library, twice as many actives or non-cytotoxic compounds are expected to be found compared to random selection. The dashed black line represents no lift.

Using the trained ML models, ∼50,000 compounds were selected from Enamine’s HTS collection (approximately 8,000,000 compounds). This included approximately 45,000 compounds in pre-plated libraries and 5,000 CP compounds. The pre-plated libraries were selected jointly by a team of medicinal chemists and a team of ML scientists based on their overall ML-predicted hit rate, predicted rate of non-cytotoxic compounds, dissimilarity to known antibiotics and compounds screened in the first stage, drug-likeness, and ADMET properties. The CP compounds were purely ML-selected; the selection logic greedily filled a budget of 5,000 compounds maximizing their predicted activity against *E. coli* lptD^-^, simultaneously minimizing their human cell cytotoxicity, and strictly avoiding compounds structurally similar to known antibiotics and compounds screened in the first stage. The details of the selection schemes can be found in the Experimental Section.

All selected compounds and compound libraries were acquired and tested for inhibition of lptD^-^ *E. coli* in a primary screen. Primary hits underwent hit confirmation with triplicate retesting in the same assay. The resulting data allowed for evaluating the forward-looking performance of the *E. coli* activity ranking models both compound-by-compound and library-wise.

Compound-by-compound predictive power was evaluated by comparing predicted and tested activities among the compounds that were selected as part of a pre-plated screening library. The same analysis was done on the CP compound set, but that yielded low apparent performance because the CP set was already highly enriched in hits and thus lacked contrast to demonstrate preferential selection. Nevertheless, the compounds selected as part of the screening libraries demonstrated robust and significant forward-looking predictive performance across all libraries (Fig. 2).

**Figure 2.**
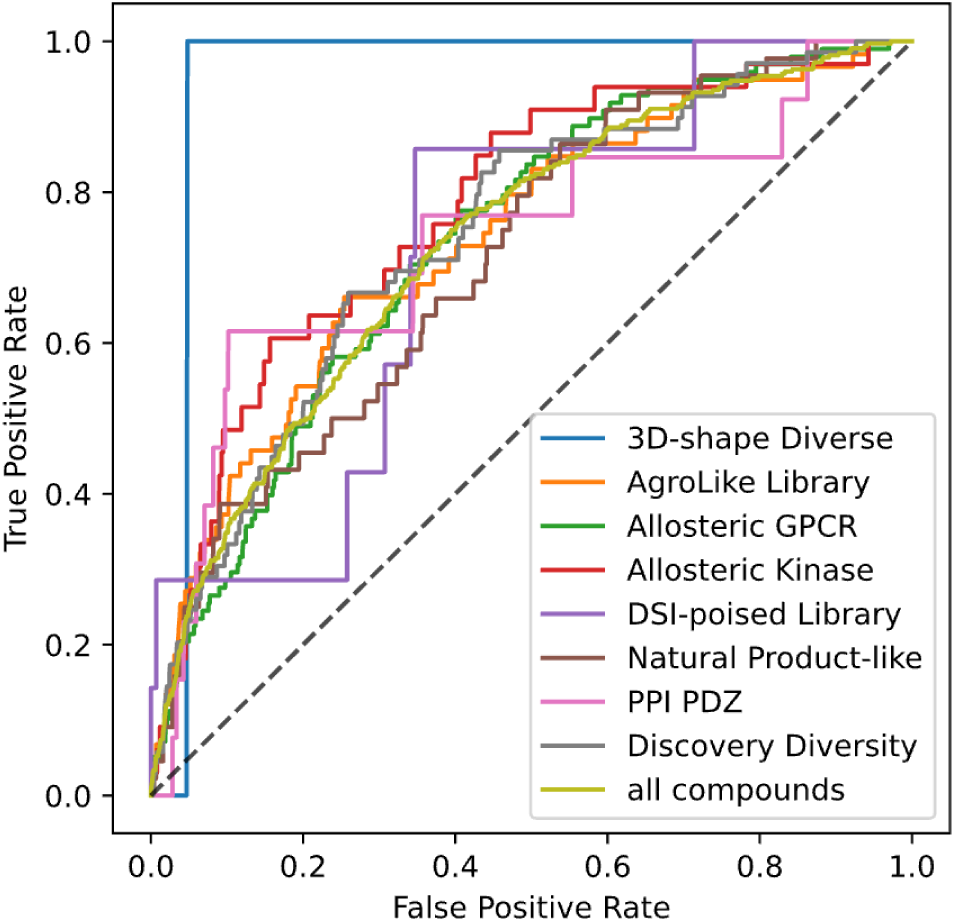
ROCs obtained by the *E. coli* ranker when the tested compound libraries are treated as forward-looking test sets. The dashed black line indicates the expectation of a random baseline.

In terms of Area Under ROC (AUROC) or lift, the forward-looking figures were approximately equivalent to the lower range of the metrics estimated in cross validation (Fig. 1): 0.73 across all tested compounds and ranging between 0.71 and 0.95 across compound libraries. This was expected for two reasons: first, backward-looking cross validation often reports optimistic estimates; second, cross validation results assumed single-objective selection, while the forward-looking results are confounded by selection for the competing non-cytotoxicity objective that is expected to lead to fewer actives. Reaching the low range of cross validation-estimated metrics even with a competing objective and dissimilarity constraints was considered a significant success and demonstrated that careful, scaffold-split cross validation estimates forward-looking performance albeit optimistically, but still within confidence bounds.

Figure 3 demonstrates the predictive power of the *E. coli* ranking model for selecting compound libraries. Each circle in the figure represents a tested compound library. The diameters of the circles are proportional to the library size. As seen, the hit rate of most compound libraries correlated strongly with their predicted quality, defined as the median predicted inhibition score. The two outliers were potentially due to either small library size (the DSI-poised library) or pronounced chemical dissimilarity to the training set (the Natural Product-like library, which contained molecules larger than training compounds). As a point of reference, we included the CP compound set in the figure as a pseudo-library. Its hit rate, 0.8%, was approximately triple the average hit rate of non-CP libraries.

**Figure 3.**
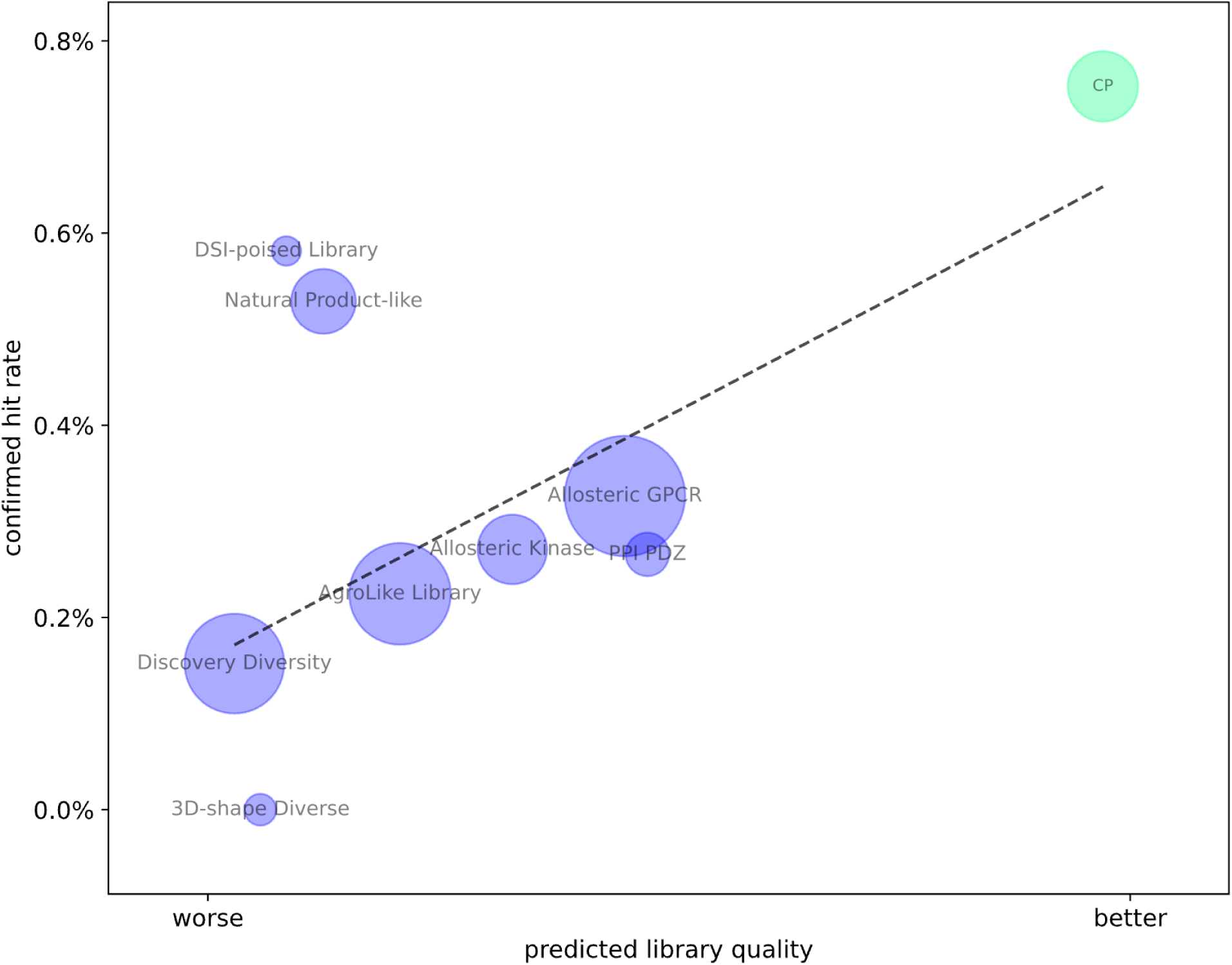
The predictive performance of the *E. coli* activity ranker in terms of predicted library quality (the median compound rank) and the confirmed library hit rate. “CP” means cherry-picked. The dashed black line is a robust linear fit.

To place our hit rates into context, we compare them with prominent literature reports: Stokes et al. reported a hit rate of 35% when testing 23 of 107 million compounds (8 out of 23 were active) (Stokes et al., 2020). Similarly, Scalia et al. reported a hit rate of 24% when testing 345 of 1.4 billion compounds (82 of the 345 were active). Both studies used an ML model trained on a curated, consistent activity dataset; besides potential structural filtering, no other ML guidance was used. At first glance, our hit rate of 0.8% seems low compared to the ones reported in the two studies; however, the following must be noted about the underlying statistical behavior:

- Selection freedom — defined as the ratio of the screening library size and the number of top predicted compounds selected — increases the apparent hit rate. In other words, when a small top fraction of a large parent library is selected, the apparent hit rates will be higher relative to selecting a larger top fraction of a smaller library.
- Optimizing for the competing cytotoxicity objective reduces the apparent hit rate.

The two references used compound-by-compound selection that more closely resembles our CP set. Here, to better compare performances, we can partially correct for the effect of selection freedom; although we cannot estimate the effect of larger parent libraries (8 million in our case vs. 107 million and 1.4 billion in Stokes’ and Scalia’s cases, respectively), we can narrow our top CP selection. When looking at the CP hit rate in the top 345 (resembling Scalia et al.), we find a hit rate of 32% (111 of 345). Similarly, when looking at the top 23 (resembling Stokes et al.), our hit rate is 48% (11 of 23). Considering that – compared to the two cited studies – our selection strategy additionally constrained for novel compounds and low cytotoxicity, we conclude that our performance is at least comparable, but likely better, than those previously reported. The likely reason for outperforming Stokes et al. is the much wider coverage of the activity model (trained on 1.2 million observations vs. 2,300 observations). It is worth noting that the extreme “elite selection” regime of the two cited studies might not be practical in a real-world drug discovery setting; since many hits are not progressible due to ADMET or other medicinal chemistry issues, most programs require, in the absolute sense, as many early hits as possible. This cannot be provided by the extremely narrow screens typically reported in the literature; therefore, the reported high hit rates may be inflated and unrealistic from a practical point of view. In our view, it makes more sense to report lifts up to a reasonable % library screened from ML-guided campaigns that explicitly consider at least a few relevant competing objectives.

The confirmed hits were counter-screened for human cell cytotoxicity. A notable improvement of cytotoxicity, both in terms of fewer cytotoxic compounds among the hits (13% in the ML-guided screen vs. 37% in the unguided screen) and widened therapeutic windows (the ratio of MICs against *E. coli* lptD^-^ and a human cell line), was observed. In addition, confirmed hits exhibited improved single-concentration inhibition compared to those of hits found in the unguided first stage of the campaign. Figure 4 shows these results.

**Figure 4.**
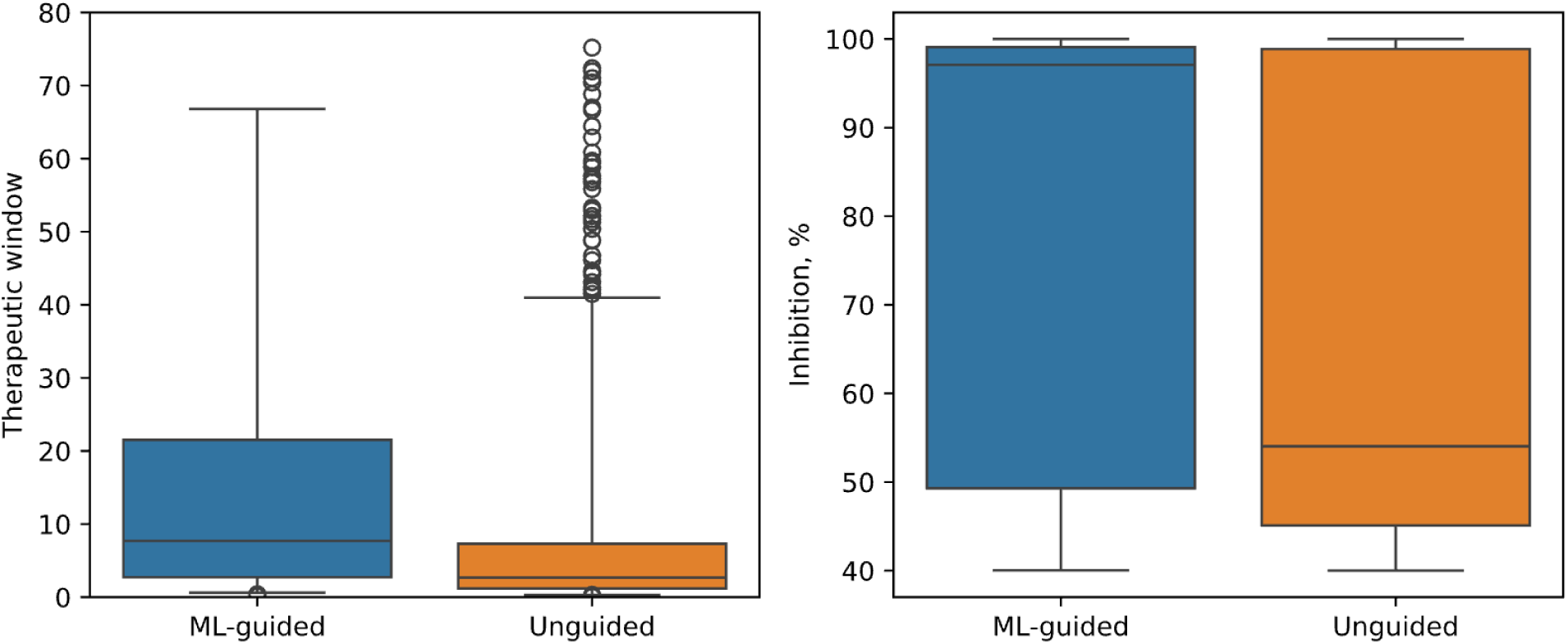
Improvement of hit quality in terms of the therapeutic window and single-point inhibition of tested compounds in the ML-guided and unguided portions of the HTS.

The structural novelty and diversity of the hits were evaluated using chemical space visualization with TMAP embedding (Probst & Reymond, 2020; Fig. 5). Visually, the TMAP embedding represents chemical space as a minimum spanning tree. Adjacent nodes represent structures that differ by a minimal structural transformation; as such, when laying the graph out using a two-dimensional force-directed layout, node distances approximate chemical dissimilarity (Kamada & Kawai, 1989). The embedding shows all screened compounds (gray), the found hits (blue), and known and approved antibiotics (red and green, respectively). As seen, the found hits covered chemical space with very few adjacent known or approved antibiotics; this implies that, on average, the found hits were structurally novel.

**Figure 5.**
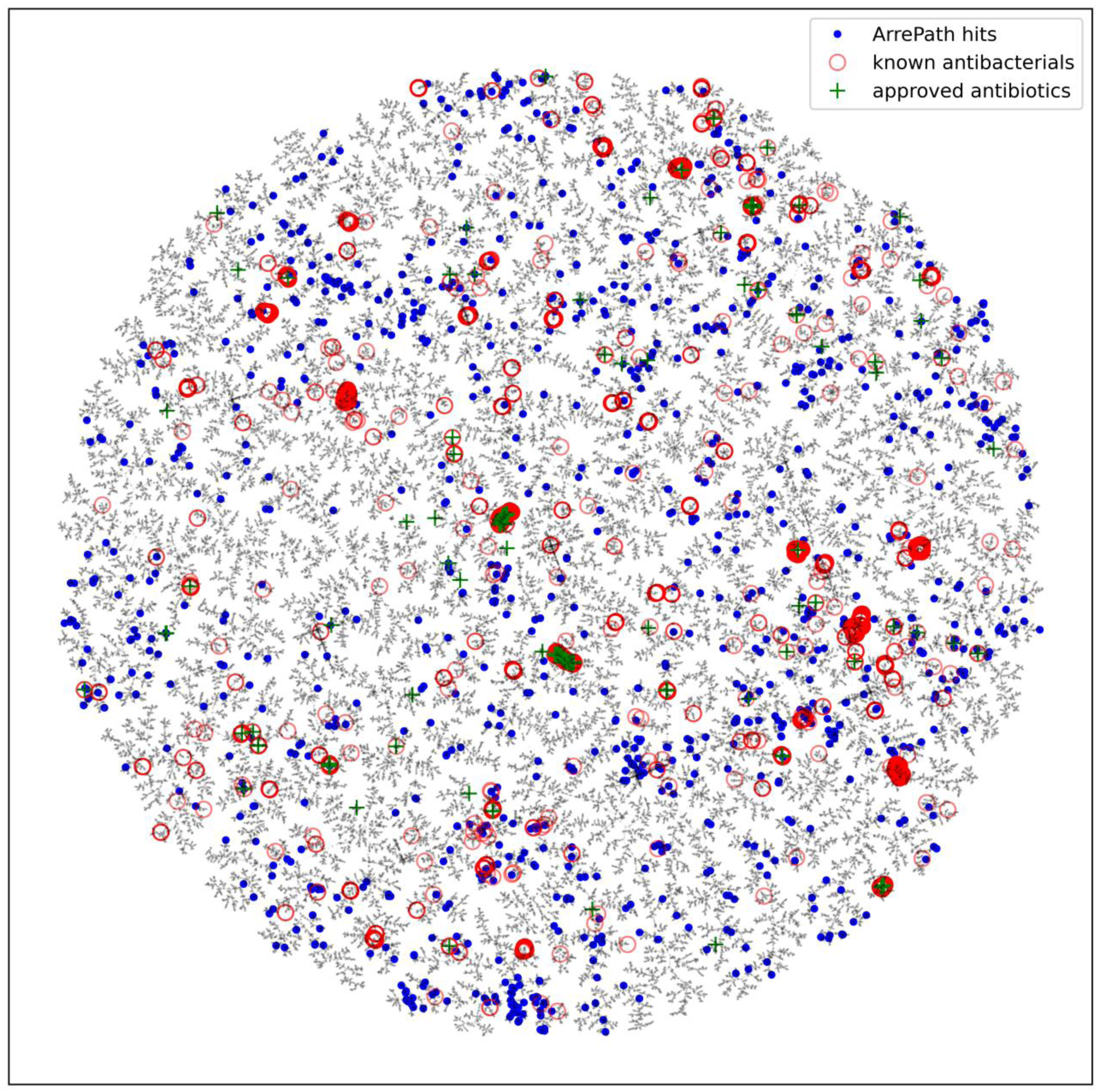
HTS chemical space in terms of its TMAP. The blue dots represent hits, the red circles represent known antibacterials, and the green crosses represent approved antibiotics. The gray dots show all screened compounds.

While most hits remain undisclosed due to their proprietary nature, Table 1 summarizes the features of five hits representing the structural novelty and drug-like features of the screening collection. All five meet Lipinski’s rule of five for oral bioavailability (Lipinski, 2004). Antibacterial potency against the screening strain on minimal media ranged from 1 to 8 μg/mL; three of the hits were also active against the parent *E. coli* strain, demonstrating the ability to permeate Gram-negative membranes. Only one, AP-00475532, had weak activity against HepG2 cells (CC_50_ = 56 μM); the rest were inactive up to 100 μM. Resistance mapping to identify the putative target/pathways of inhibition suggest diverse MOAs, including nucleotide, amino acid, vitamin or cell wall biosynthesis pathways. These pathways are only essential in minimal media; the lack of antibacterial activity of all five hits in rich media supports the putative MOAs identified by resistance mapping. The metabolic pathways these molecules target are likely non-essential in many sites of infection, making these molecules unattractive for further development. For completion, we note that while these molecules are not further pursued, two programs have resulted from hits identified in these screens: at the time of writing, one in late-stage lead optimization and one in early hit-to-lead optimization.

## CONCLUSION

A Machine Learning (ML)-guided High Throughput Screening (HTS) campaign for finding small molecules that inhibit the growth of *E. coli* was carried out in a commercial setting. The HTS was guided by Learning-to-Rank (L2R) ML models that predicted growth inhibition and human cell cytotoxicity. The L2R approach allowed training on uniquely large training datasets. 50,000 compounds were acquired: 45,000 in pre-plated libraries and 5,000 individually selected. The compounds were tested for *E. coli* growth inhibition. The confirmed hits were counter-screened for human cell cytotoxicity. In the individually selected compounds, ML-guiding doubled the hit rate and reduced the number of cytotoxic compounds to one-third relative to a human medicinal chemistry baseline, yielding an overall tripling of progressible hits. Among the very top compounds, the observed hit rate was 32-48%, in line with other prospective ML-guided antibiotics HTS studies, except that we achieved this hit rate while optimizing for the competing low-cytotoxicity objective and explicitly screening for structurally novel compounds. The results highlight the potential of ML guidance in early-stage drug discovery.

## EXPERIMENTAL SECTION

### Learning-to-Rank for antibiotics discovery

An important concept in ML is generalization (extrapolation), which, in the drug discovery domain, translates to retained predictive power over chemical dissimilarity. In other words, an ML model that generalizes remains accurate when predicting properties of compounds that are dissimilar to compounds in the training set. Naturally, generalization is critical in drug discovery since only novel compounds can form intellectual property; furthermore, chemical novelty is linked to MOA novelty, which in turn is highly valuable from the point of overcoming resistance to antibiotics. ML models are known to extrapolate beyond their training data up to a certain structural dissimilarity. While modern model formulations generalize better than conventional QSAR models, this effect is of lesser magnitude compared to the effect of expanded chemical space coverage due to a larger and more diverse training set.

The main idea behind our ML modeling approach is that measurement noise in HTS datasets dominates prediction performance over model complexity, making broad chemical space coverage more valuable than complex model formulations. This leads to the question: how do we increase chemical coverage?

Conventionally, ML models in drug discovery are trained on curated datasets that only contain observations consistent in terms of assay conditions. High-quality, consistent, and curated datasets are often cited as important pre-requirements of ML-guided drug discovery. The public space is ripe with high-quality datasets that could expand the coverage of training data; however, these HTS datasets are often acquired at different experimental conditions and are thus incomparable to each other and to proprietary data. The consolidation of disparate data sources is therefore important to improve ML performance.

Data consolidation can occur in two ways: data-based and model-based. Data-based consolidation is data curation and transformation; for example, HTS datasets can be merged by carefully extrapolating inhibition values to higher or lower drug concentrations. Although these transformations are more explainable in terms of the underlying science, they are time-consuming and often limited (Landrum & Riniker, 2024). In contrast, the model-based approach elects model formulations that are insensitive to inconsistencies between datasets and can utilize multiple disparate sources optimally, without the need for significant a priori data transformation and curation.

From the ML literature, we mention two routes that yield models that can utilize multiple data sources: multitask learning and Learning-to-Rank (L2R). Multitask learning uses model formulations that allow for learning from multiple datasets; multitask models were reported as promising alternatives to single-task models in drug discovery (Capela et al., 2019; Ramsundar et al., 2015; Xu et al., 2017). Typically used to improve models by jointly learning to predict multiple, correlated target properties, the main drawback of the multitask formulation is having to train prediction “head” models for each prediction task. While, when properly parametrized, the head models can harmonize multiple tasks, the resulting multitask model might lack a coherent scoring function that can prioritize compounds by a single property. Besides classifiers and regressors, L2R is a major type of ML approach that is concerned with ranking or prioritizing queries based on their relevance (Cao et al., 2007; T.-Y. Liu, 2009). Although L2R was evaluated for drug discovery and molecular property prediction (Agarwal et al., 2010; Rathke et al., 2011; Zhang et al., 2015), it has not been used in HTS campaigns before. The simplest form of L2R learns to compare pairs of observations and predict whether the first or second in each pair is of higher “relevance” – relevance in a drug discovery application might refer to inhibition, binding, or any other property to be optimized; for example, antibiotic compounds that are predicted to be more active are assigned higher relevance. When applied to prioritizing compounds, L2R exploits the approximate invariance of compound ranks to assay conditions; for example, within reasonable bounds, it is expected that the relative relevance of a pair of compounds observed in a WT assay is similar to that observed in an assay of a mutant strain. As such, when learning relative compound relevance, an L2R model can learn from multiple, disparate datasets (see Fig. 6). The heterogeneous data can come from different HTS assays, can be reported in terms of different endpoints (e.g., a single-concentration inhibition or an MIC value), and can overlap if the underlying information (compound relevance) is approximately consistent. In practice, compound relevance is a continuous scalar; when unseen compounds are sorted by their predicted relevance, they will be approximately ordered by the target property, e.g., inhibition. We trained two L2R models: one for ranking by *E. coli* lptD-activity and one for ranking by human cell cytotoxicity. The L2R formulation allowed us to utilize multiple sources in the training data resulting in models with state-of-the-art chemical space coverage: the *E. coli* lptD-activity ranker was trained on over 1.2 million observations of approximately 700,000 unique compounds, while the cytotoxicity ranker was trained on over 2 million observations of 520,000 unique compounds. Table 3 is a summary of training data sources.

**Figure 6.**
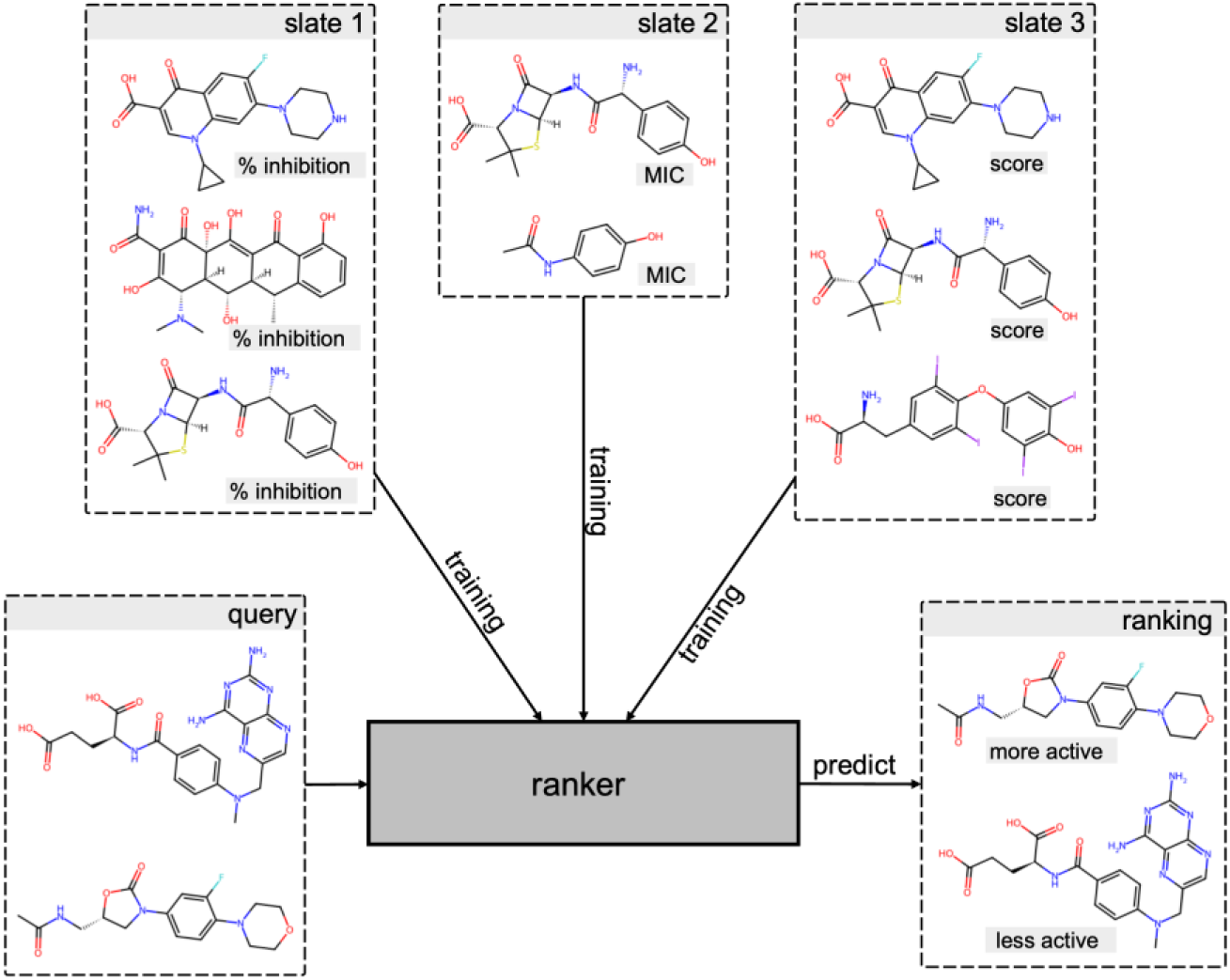
L2R concept. During training, an L2R model learns to compare pairs of compounds, based on e.g., their relative growth inhibition. During prediction, the model outputs a single score; higher scores indicate higher compound relevance, i.e., higher predicted inhibition. The score can be used to, e.g., rank-order a virtual library.

**Table 3.** Data sources and dataset sizes for *E. coli* inhibition and human cell cytotoxicity training data. The numbers are the numbers of experimental observations in each dataset.

| Data source | *E. coli* inhibition | Cytotoxicity |
| --- | --- | --- |
| Proprietary | 400,000 |  |
| Related patents | 140 |  |
| CO-ADD | 225,000 | 4,600 |
| PubChem | 490,000 | 2,114,000 |
| ChEMBL | 6,600 |  |

### Compound-by-compound selection

The CP selection scheme was designed to select compounds one-by-one from Enamine’s Liquid Stock collection while maximizing the number of hits, minimizing the number of cytotoxic compounds, and simultaneously avoiding known antibiotics and other non-novel compounds. The scheme comprised preferential selection and constraints. For simplicity, we discuss the constraints first. Using Extended Connectivity Fingerprints (ECFP), a Tanimoto distance cutoff of 0.6 was used to exclude compounds similar to known or approved antibiotics and compounds screened in the first stage of the HTS campaign. The remaining chemical space was scored by the two ranking models to predict compound relevance in terms of *E. coli* activity and non-cytotoxicity. Figure 7 selection shows a two-dimensional histogram of the achieved scores. The inverse correlation between inhibition and cytotoxicity was immediately apparent. This was expected, as non-specific toxicity is a prevalent factor in bacterial inhibition; therefore, the selection logic had to realize optimization over two competing objectives. To identify an optimal tradeoff between activity and non-cytotoxicity, a small, held-out, independent dataset was used to estimate the inverse linear relationship between the two objectives using a Logistic Regression (LR) model. Starting from the non-cytotoxic and active (top right) quadrant in Fig. 7, the normal of the LR decision function was used as the direction of greedy compound selection. Compounds that achieved the best aggregate LR score were greedily picked. The chemical space was sphere-exclusion clustered and the cluster neighbors of every picked compound were excluded; therefore, only the best cluster member was picked per next-best compound. Additionally, an arbitrary activity threshold of the 65^th^ percentile of the predicted activity scores was used. Selection proceeded this way iteratively, picking the next-best compound in terms of its aggregate score, and excluding its cluster members from further selection until the quota of 5,000 diverse, novel compounds was filled. Note that the zero isoline of the LR model was not approached, indicating that the parent library had remaining potential for further selection. The selection frontier in terms of the aggregate LR score is denoted as “decision margin” in Fig. 7.

**Figure 7.**
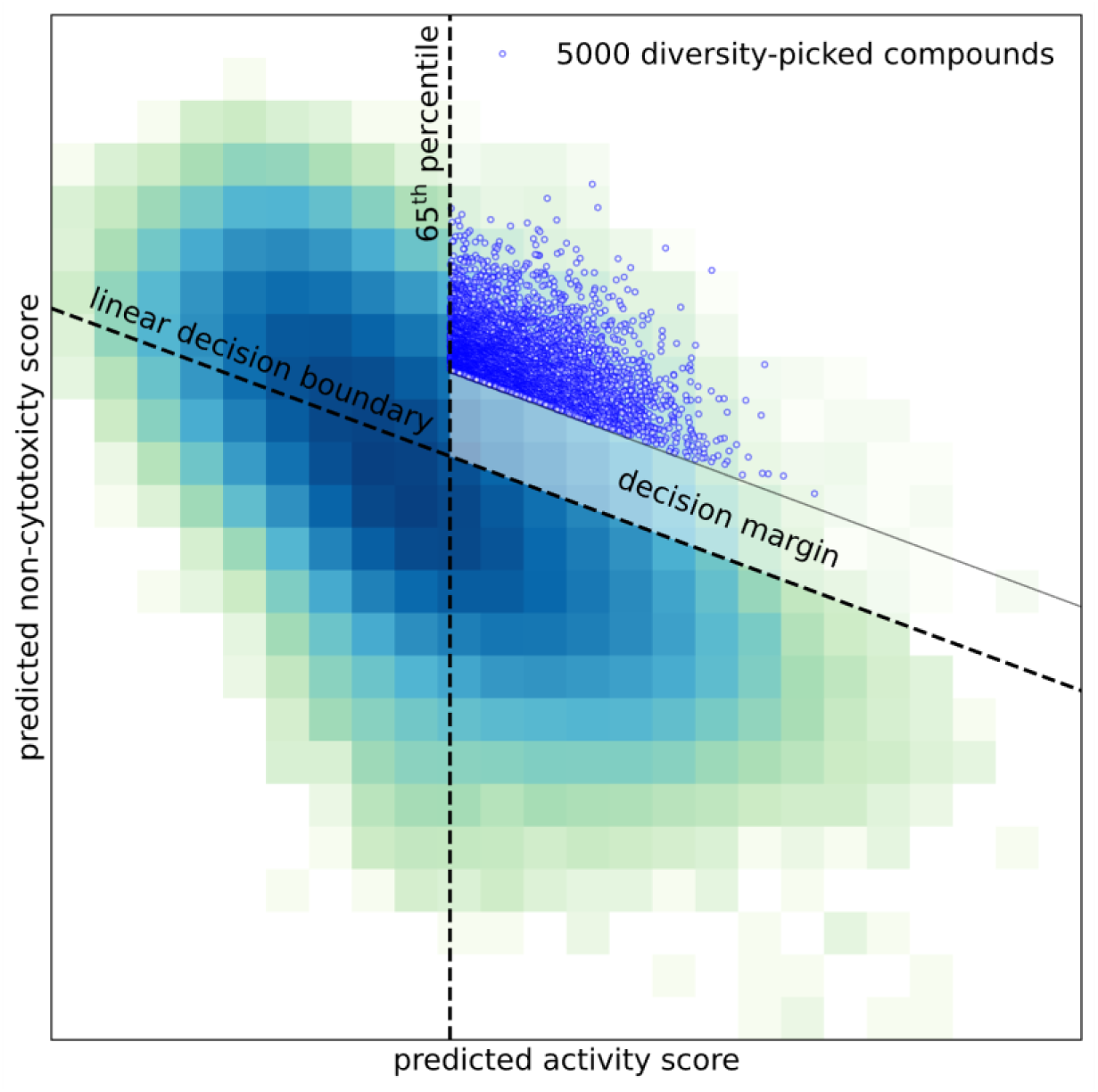
Cherry-picking selection logic. Blue hues show a 2D histogram of compounds in the space of predicted non-cytotoxicity and activity (dark blue indicates high density; high non-cytotoxicity score implies a high likelihood of non-toxicity; high activity score implies high likelihood of activity). The inverse correlation signifies that generically toxic compounds are more likely to be antibiotic. The linear decision boundary is obtained as the separating line of a logistic regression model that best separates cytotoxic and not cytotoxic compounds in an independent test set. The decision margin represents the boundary that encompasses the cherry-picked compounds by greedily filling a selection budget optimizing the predicted therapeutic window and compound diversity. In addition, compounds that were predicted to be less active than the 65^th^ percentile of predicted activity were excluded.

### Microbiology

High throughput screening (HTS) was conducted at Evotec (Toulouse, France). The screening collection was tested for antibacterial activity at 30 μM against 5 x 10^5^ CFU membrane-permeabilized *E. coli* lptD4213 (Sampson et al., 1989), which was incubated for 24 h at 37 °C in in 30 μl M9 minimal media (+2% glucose) in a 384 well plate. After equilibrating the plate to room temperate, 10 ul of Bac TiterGlo^TM^ reagent (Promega cat # G8233, prepared according to the manufacturer’s instructions) was added, mixed briefly on the orbital shaker, incubated for 5 min and luminescence was recorded with an integration time of 100 ms. Vancomycin and tetracycline were used as positive controls. Hit confirmation was done in a triplicate single concentration retest at 30 μM. Hit profiling was performed in duplicate in an 11-point dose response assay (1:2 dilutions from a top concentration of 100 μM). Cytotoxicity testing was also performed at Evotec in 384 well plates, with each well containing 3,000 HepG2 cells (ATCC cat # HB-8065) in 20 μl culture media (Dulbecco’s Modified Eagle’s Medium [DMEM, Gibco cat # 11885-084] supplemented with 10% fetal bovine serum [FBS, Gibco cat # A5669701], 1x penicillin-streptomycin mixture [Solarbio cat # P1400], 1x non-essential amino acids [NEAA, Gibco cat # 11140-050] and 1% HEPES [Gibco cat # 15630-080]) incubated for 48 h at 37°C, 5% CO_2_ with test compounds in the same dose response set up as above. After incubation, 20 μl of CellTiterGlo reagent (Promega, cat # G7571), prepared according to the manufacturer’s instructions) was added. Plates were incubated at room temperature for 10 min, then luminescence was measured as described above. Resistance mapping was performed by plating up to ^10⁹^ CFU *E. coli* lptD4213 on M9 minimal media (+2% glucose) agar at 4X the agar MIC of selected compounds, incubated up to 48 h at 37°C. Once stable resistance of isolated colonies was confirmed by MIC testing, resistance mapping was performed by comparing genomic sequence of mutants to the parent strain (sequencing was performed at SeqCenter [Pittsburgh, PA] and analysis was performed using breseq software (Deatherage & Barrick, 2014)).

## AUTHOR INFORMATION

### Author Contributions

The manuscript was written through contributions of all authors. All authors have given approval to the final version of the manuscript.

## ABBREVIATIONS

ML: Machine Learning
CP: cherry-picked
L2R: Learning-to-Rank
ROC: Receiver Operating Characteristic
AUROC: Area Under Receiver Operating Characteristic Curve.

